# Does field margin semi-natural habitat amplify the abundance of beneficial spiders in a vineyard canopy?

**DOI:** 10.1101/2025.07.30.667680

**Authors:** Cord Phelps

## Abstract

I compared spider counts from a transect bordered by a semi-natural habitat to those of a control transect in a California Central Coast organic vineyard over the course of the 2018 growing season, observing spiders in blue vane traps that had been suspended in the canopy fruit zone. Ranked abundance analysis of spider counts did not confirm a difference between the two transect populations for any time period and a population gradient proceeding from the vineyard edge into the vineyard center was not identified for either transect. I suggest that semi-natural habitat proximity does not promote spider populations in the ecological context of a classical Central Coast vineyard.

## 1 Introduction

Semi-natural habitats (SNH) are believed to contribute various ecosystem services in agricultural settings. Faced with an array pest management challenges, the organic wine grape industry and associated research institutions have evaluated the potential contribution of SNH to biological control strategies. However, conclusions are often contradictory and qualitative. This study evaluates the sensitivity of beneficial spider populations to the proximity of SNH in a California Central Coast organic vineyard. The data suggests that there is no effect.

While vineyard biodiversity amplifies arthropod predation and parasitization (Altieri et al. 2010), both semi-natural habitats and cover crops have been studied for many years as a refuge for, and source of, insects that antagonize vineyard pests. However, the climate of the Central Coast and general scarcity of irrigation water limits the persistence and utility of cover crops as a driver of biodiversity. As a consequence, more specific quantification of SNH pest regulation contributions and simplified, grower friendly, predator observation tools are of interest.

This study was designed to identify beneficial insects that can be easily observed by Central Coast grape growers and to determine if SNH appears to promote beneficial arthropod populations in an organic vineyard.

## 2 Materials and Methods

### 2.1 Study Site

#### 2.1.1 Ampelos vineyard, Lompoc, California

My 2018 study was hosted by Ampelos Cellars in the Sta. Rita Hills American Viticultural Area (AVA) of Santa Barbara County, California. The growing season of this AVA is characterized by heavy marine fog in the mornings and clearing with a steady wind in the afternoon. (Appendix B)

I selected a control transect bordered by a plowed field and a SNH transect with several large native oak trees at its base. These two transects have a matching orientation to the prevailing wind and are 300 feet apart. Vertical shoot positioned (VSP) Pinot Noir is planted at 4 foot intervals in each transect. Vineyard rows are at a 10 foot spacing. Native ground cover was not maintained.

### 2.2 Sampling Procedures

The two transects, each populated by 30 blue vane traps with yellow jars (BanfieldBio 2025), were sampled twice per day, for 12 weeks during the growing season. Traps were suspended in the canopy fruit zone connected to a VSP canopy positioning wire.

The 30 traps were positioned in 3 vineyard rows of 10 traps; 6 traps at 16 foot spacing (at 4, 16, 32, 48, 64, and 80 feet from the vineyard edge), and 4 traps at 100, 130, 160 and 200 feet. Rows were separated by an un-sampled row.

Each vane trap was inspected at 0700 and 1700, the species that were present inside the vane trap bowl were recorded and released. The trap was then re-assembled and replaced in its previous position.

At the end of a typical 3 day sampling week, the traps were shifted to an adjacent row (un-sampled in the prior week).

### 2.3 Analysis of Collections

Count data for Thomisidae, the most easily recognized numerous beneficial acquired in the study, is non-normal, and zero-inflated. Analytical methods were selected to compare distributions between transects (control vs. SNH) and among transects (row to row). Spatial and temporal effects were also evaluated.

#### 2.3.1 Non-parametric methods

Mann-Whitney U Test (Wilcoxon Rank Sum Test) is used to compare the two independent transect groups to assess differences in their distributions. (Valentini et al. 2022; James, Link, and Pyle 2013)

The Wilcoxon Signed Rank Test is used to compare two matched samples of the same-transect row data to assess similarity based on median difference. (Nicholls, Parrella, and Altieri 2001)

#### 2.3.2 Resampling Methods

To double-check classical transect comparison result, a resampling generative model was used to accommodate zero-inflated count data. Simulated data from a binomial distribution (1000x) with a parameter representing the probability of sampling Thomisidae at a single trap position results in credible intervals of population density. (Seavy et al. 2005).

Results were assessed using bayestestR, with a focus on comparing the credible intervals of posterior distributions (Makowski, Ben-Shachar, and Lüdecke 2019; Inácio and Rodríguez-Álvarez 2023)

#### 2.3.3 Presence/Absence methods

Presence/absence similarity of same-day, same-time, same-transect row samples is evaluated using the Sørensen Index (Hogg and Daane 2011).

Binomial proportions and confidence intervals of an individual trap collecting a Thomisidae specimen in any one week period is calculated by the exact binomial test of Clopper-Pearson (Oregon Wine Research Institute 2018), binom.test (R Core Team 2025). These CIs are known to be conservative and useful when the sample size is < 30.

#### 2.3.4 Distance effects

Spatial association as expressed by population clusters as a function of trap position was assessed using Bray-Curtis similarity (Zhao et al. 2023; Ricotta and Podani 2017; Zarraonaindia et al. 2018)

Spatial dissimilarity analyses were performed in R (version 4.3.0; R Core Team, 2023) using the vegdist function from the vegan package (version 2.6-4; Oksanen et al., 2022) and the constr.hclust function from the adespatial package (Dray et al., 2022).

#### 2.3.5 Original methods

Binomial patterns of spatial Thomisidae presence were interpreted as N-Grams and used to score ‘similarity’ between same-day, same-daytime, same-transect row pairs. These results are used to assess the sensitivity of the calculated Sørensen Index.

## 3 Results

3,720 trap observations were made between June and August with 4,583 arthropods recorded and released.

Thomisidae was a numerically dominant predator that can be easily identified by an untrained entomologist. (Appendix A)

### 3.1 Inter-transect similarity

The Mann-Whitney U test results indicate that the data does not contain enough evidence to suggest that Thomisidae count data for the SNH and control transects originate from different distributions. (minimum reported p-value = 0.13).

Re-sampling methods found that the overlap coefficient of the SNH and control Thomisidae population posterior distribution indicates ambiguous divergence and moderate similarity of the first two sampling periods.

The Thomisidae population had largely disappeared from the canopy by week 32.

## 4 Discussion

“SNH” is characterized by reduced management activity and the presence of native vegetation (Mestre et al. 2018). On the Central Coast, vineyards are often surrounded by a SNH profile composed of grasslands used for grazing interspersed with oak trees (*Quercus douglasii*).

The crab spider (*Thomisidae*) is a predator of common vineyard pests including leafhoppers (*E. elegantula*) (Altieri et al. 2005), (Costello and Daane 1999). Using the blue vane trap as a sampling tool, crab spiders were easily collected.

Prior research suggests that insect density decreases moving from the vineyard edge ((Thomson and Hoffmann 2009)), I had hypothesized counts would rise to some relative peak near the vineyard edge and then gradually decline. The blue vane trap layout and sampling strategy was intended to quantify that effect in this specific vineyard layout and management protocol.

### 4.1 Interception of arthropods with vane traps

The grower asked that I not introduce any chemicals as part of the study or use sampling techniques that would remove insects from the vineyard population. Consequently, the more common techniques of using sticky traps or pitfall traps were not available for this study.

Blue vane traps seemed to be the only viable option for collecting and releasing insects generally. Vane traps have been used to sample Hymenoptera in other studies (Kimoto et al. 2012; Hall 2018) but apparently not for Araneae and never suspended in vineyard canopies.

Thomisidae readily presented themselves in the bowls of the vane traps and were the most frequently encountered beneficial arthropods (Appendix A). However, Thomisidae counts of ‘0’ were often recorded. I did not attempt to estimate the vine-local Thomisidae population, and therefore am uncertain about the efficiency of using a vane trap to sample these arthropods.

#### 4.1.1 Diversity of sampled insects

#### 4.1.2 Thomisidae sampled population overview: SNH and control transects

The morning sample usually represented animals that encountered the trap in the 12 to 14 hours since the afternoon sample of the previous day. These groups were numerically similar. Without a better understanding of the prey and predator vineyard ecology, I elect to analyze these two groups separately even though there is some prior research that suggests they could be combined. (Costello and Daane 2005)

#### 4.1.3 Temporal Thomisidae distribution

Thomisidae presence was more pronounced in June (weeks 23-25) than it was in August (weeks 32-34).

#### 4.1.4 Spatial Thomisidae distribution

Weekly counts were zero-inflated but the average count per trap was relatively constant although the control transect exhibits a slight count gradient as traps are positioned deeper into the vineyard rows.

Local micro-clusters of Thomisidae seemed to ‘float’ among adjacent trap positions from week-to week, Figure 7 provides an example from weeks 25 and 30. Neither transect exhibited ‘uniform’ lateral patterns of daily Thomisidae counts in the 3 adjacent rows, nor did the SNH population seem to establish a predictable gradient of counts proceeding from the vineyard edge into the vineyard center. The existing research discussing Thomisidae ballooning led me to expect to observe a count maximum close the the SNH vineyard edge followed by a decline to an approximate vineyard average Thomisidae presence (Holland, Perry, and Winder 1999), (Pearce and Zalucki 2006).

**Figure 1.**
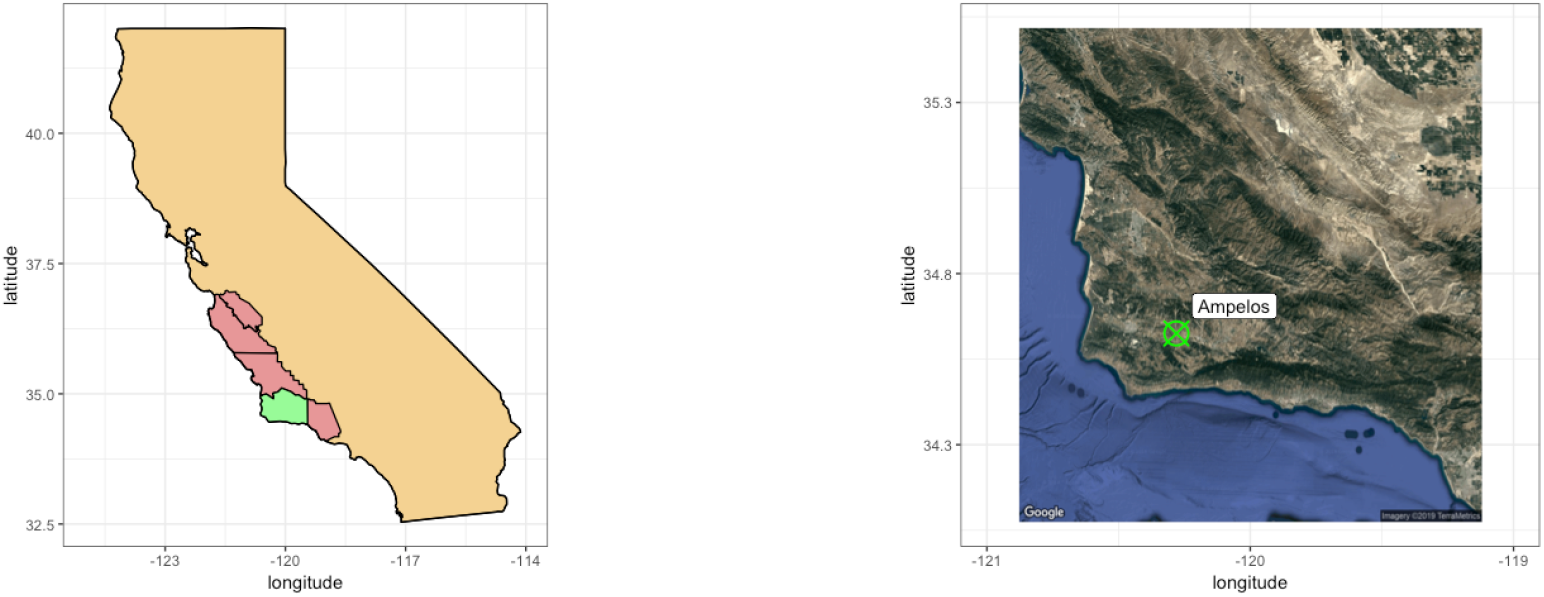
Central Coast California.

**Figure 2.**
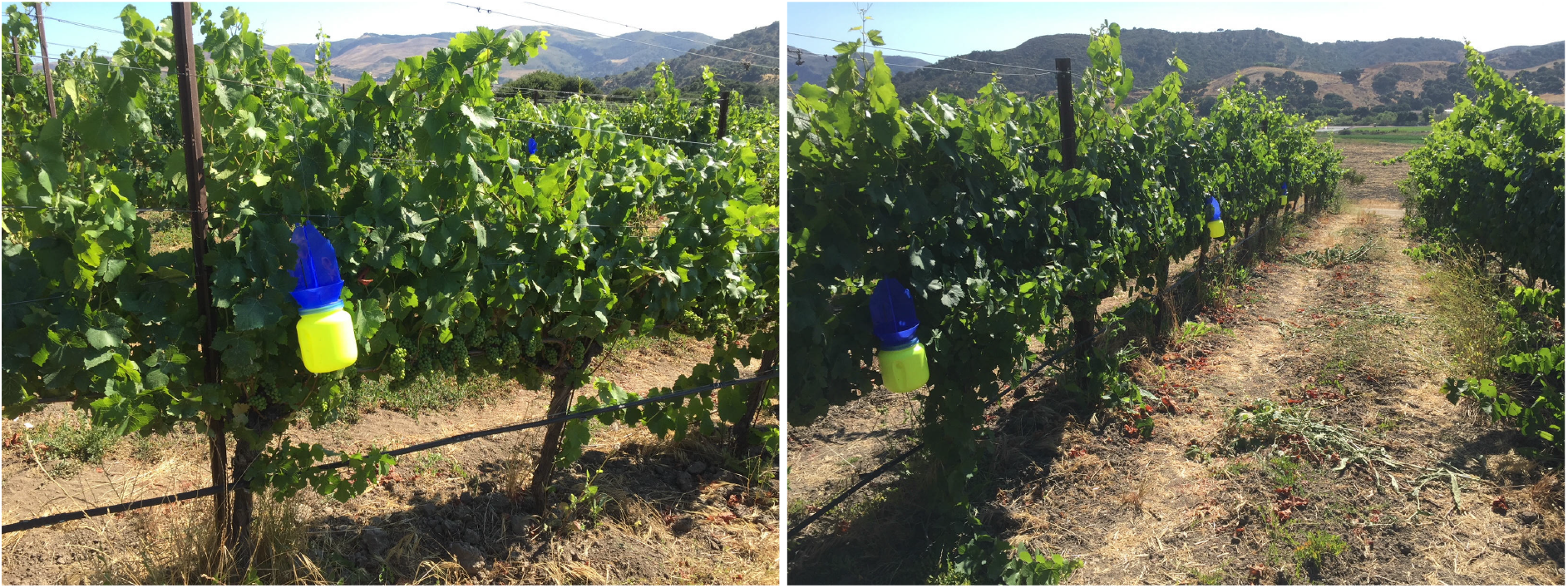
Vane traps in the canopy fruit zone.

**Figure 3.**
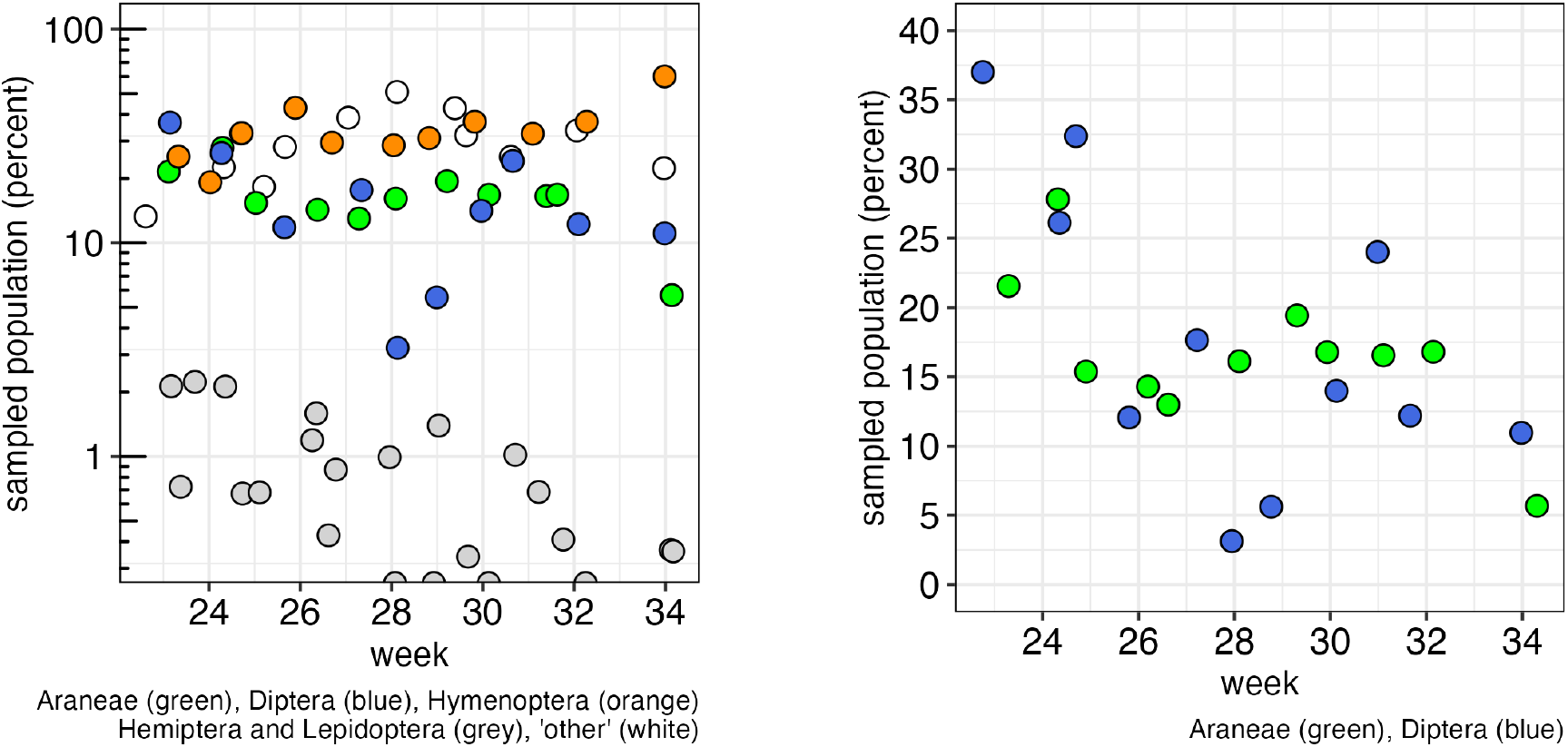
insect abundance by taxonometric Order.

**Figure 4.**
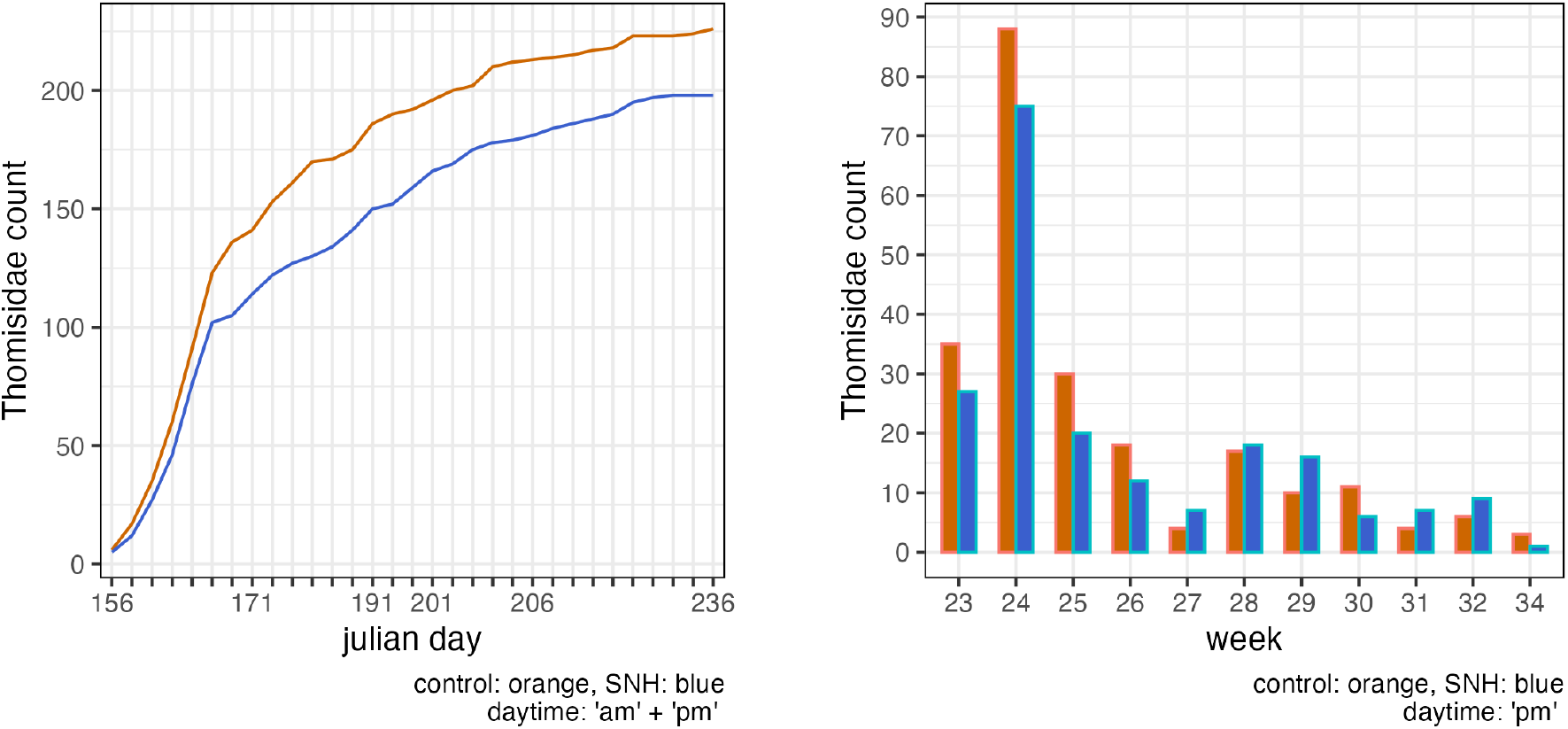
Cumulative Thomisidae counts.

**Figure 5.**
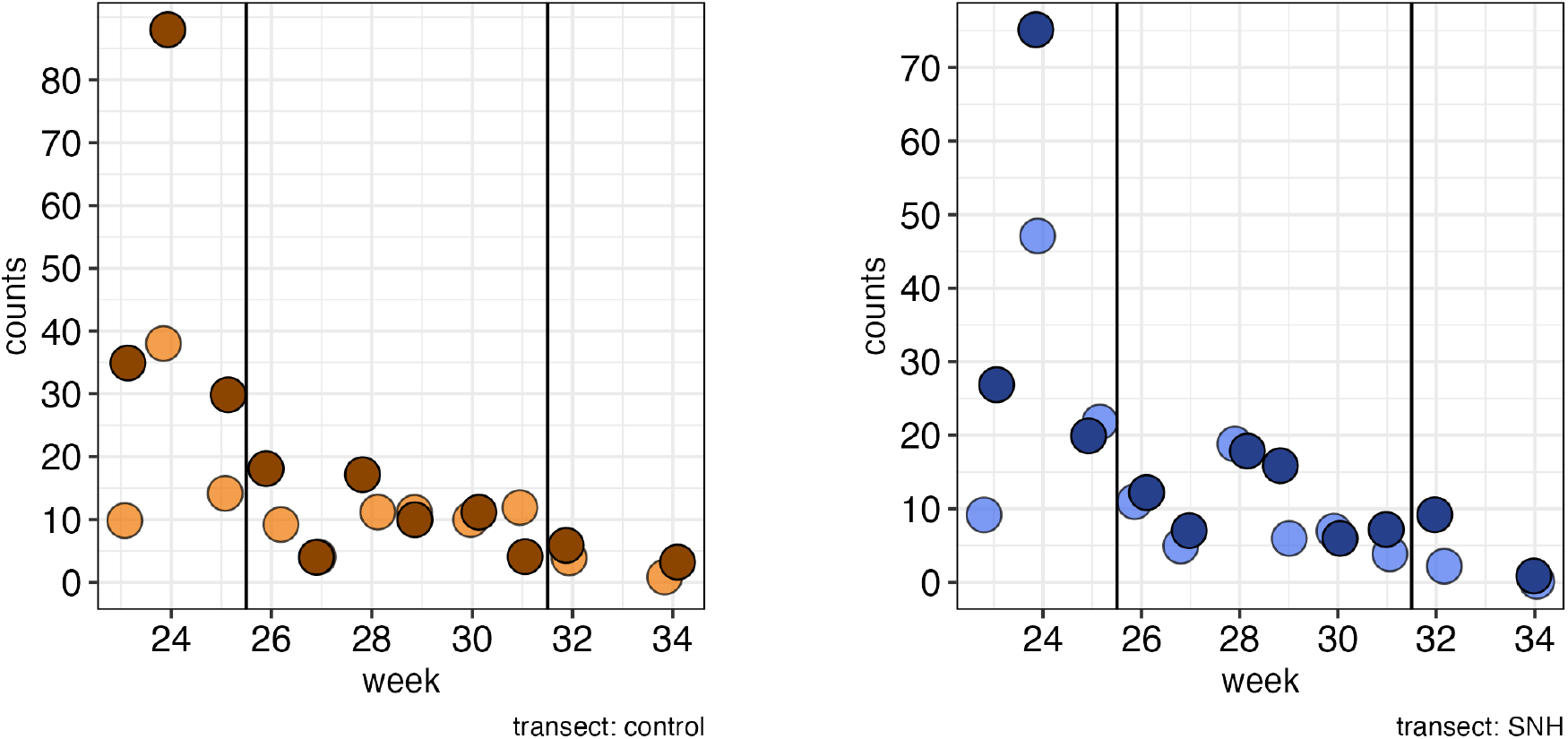
Weekly Thomisidae counts and 3 proposed temporal clusters: weeks 23-25, weeks 26-31, weeks 32-34 (“pm”: dark color, “am”: light)

**Figure 6.**
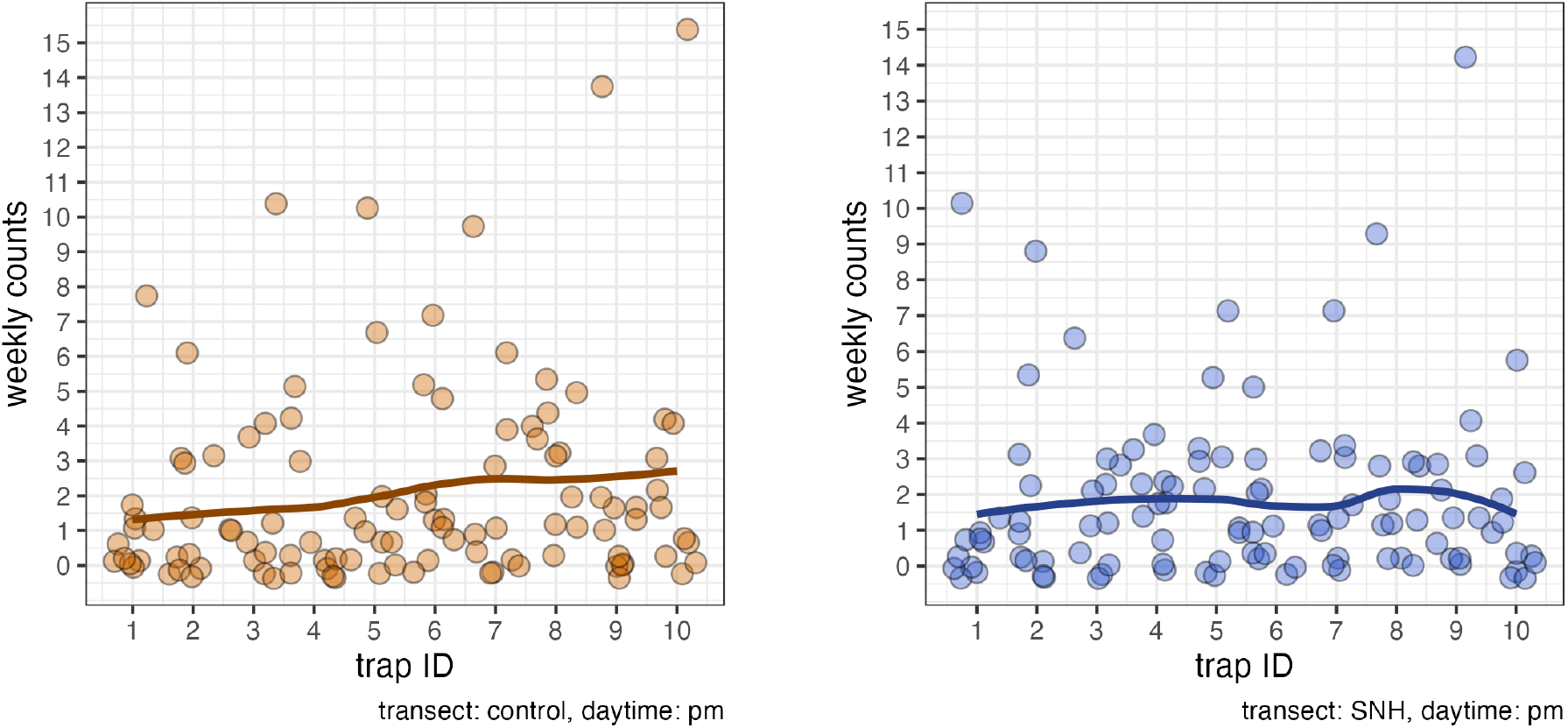
Thomisidae weekly counts (jittered) and average (solid line) by trap ID.

**Figure 7.**
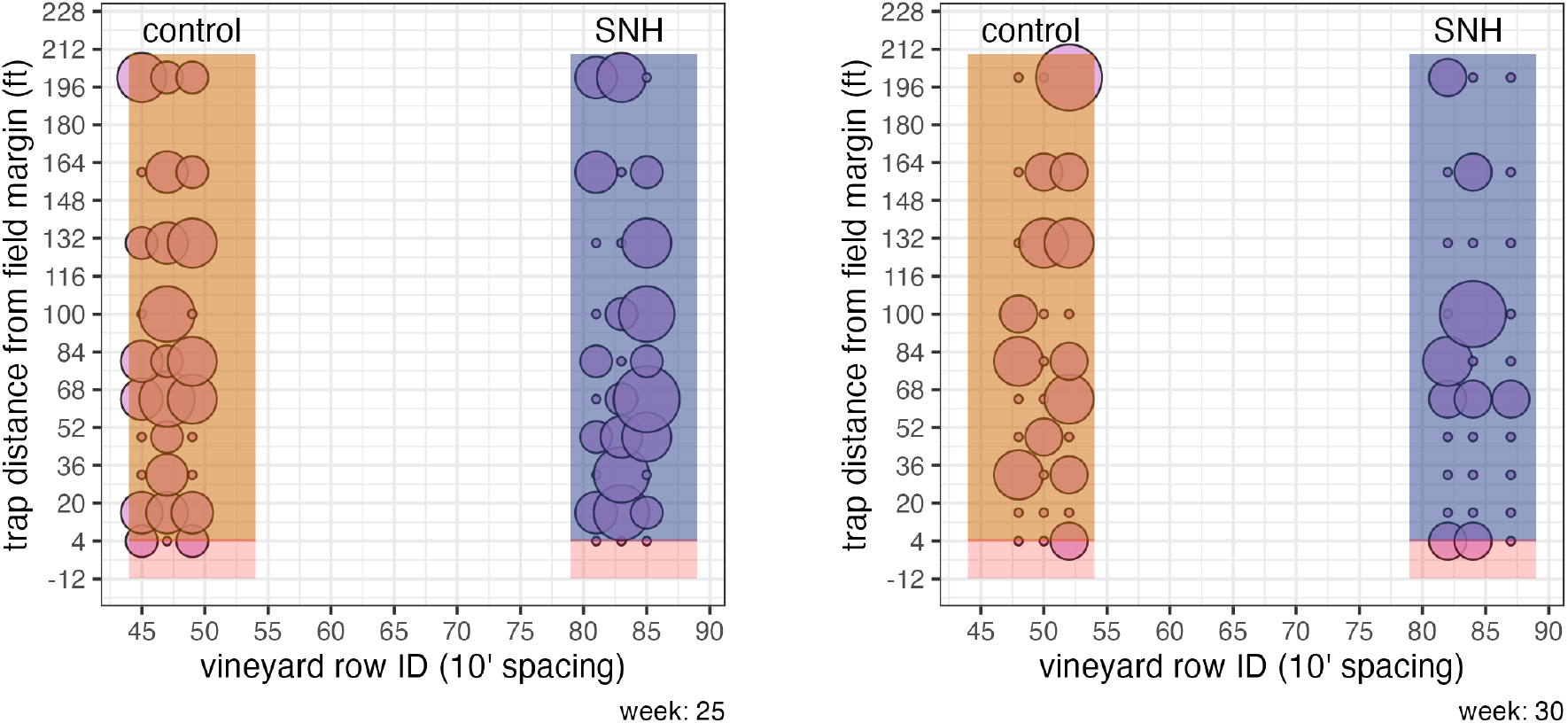
Thomisidae presence: weeks 25 and 30.

##### Spatial population clusters

I imagined that the population would increase suddenly from the SNH vineyard edge and then decrease gradually as sampling progressed into the vineyard.

ANOVA has been used to determine the influence SNH relative to the vineyard edge (Thomson and Hoffmann 2009), but this study could not satisfy the ANOVA expectation of normality. As such, I imagined that at least 3 population clusters might be identifiable. I used spatially constrained hierarchical clustering to describe 3 weekly clusters in each transect with clusters formed based on Bray-Curtis similarity to adjacent positions (Zhao et al. 2023).

##### Bray-Curtis dissimilarity matrix

The Bray-Curtis Dissimilarity Matrix was computed using the vegdist function from R library vegan (Dray et al. 2025). Counts were aggregated, but not transformed.

Spatially constrained clustering was computed using the constr.hclust function from R library adespatial (Guénard and Legendre 2022) and the result for both transects suggests a population transition at about trap position 3 and another population domain in the trap position range 7 to 10 (Figure 8).

**Figure 8.**
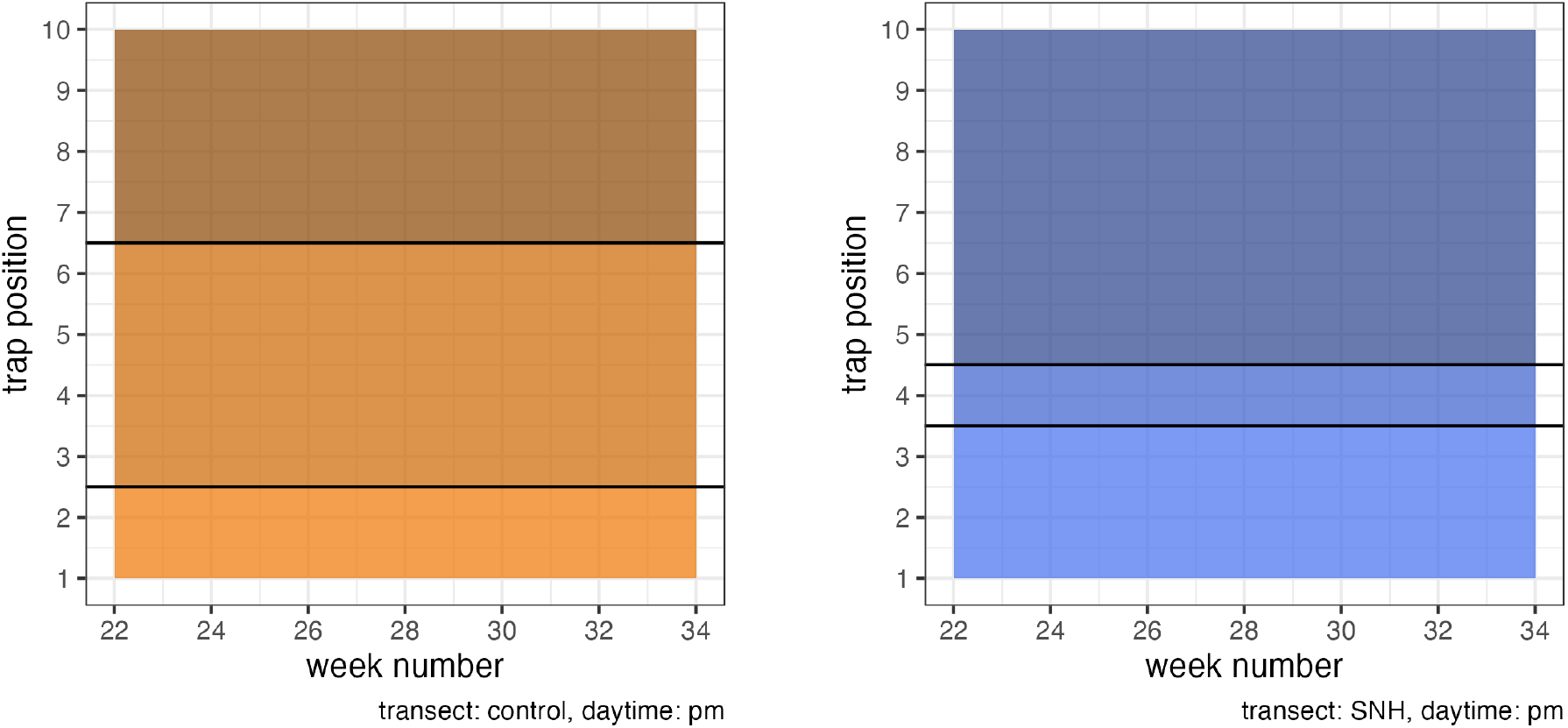
Constrained hierarchical clustering of weekly Thomisidae counts.

Bray-Curtis (Figure 8) visually suggests that the first three trap positions are recording a population that is different from the last three trap positions. However, the Mann-Whitney U test does not confirm that result. The assessment is that the data does not contain enough evidence to conclude that the intra-transect comparison of clustered counts of traps 1-3 and clustered counts of traps 8-10 come from different population distributions both for the control transect and for the SNH transect. (Table 3)

**Table 1:** Thomisidae counts, SNH vs control, daytime=‘pm’ Mann-Whitney U Test (Wilcoxon Rank Sum Test)

| timeframe | sampling time | U statistic | p-value |
| --- | --- | --- | --- |
| weeks<br>23 - 25 | ‘pm’ | 70.5 | 0.13 |
| weeks<br>26 - 31 | ‘pm’ | 42.5 | 0.59 |
| weeks<br>32 - 34 | ‘pm’ | 48.0 | 0.90 |

**Table 2:** Similarity assessment of SNH and control posterior count distributions (bayestestR::overlap)

| timeframe | sampling<br>time | Overlap<br>Coefficient | similarity<br>assessment |
| --- | --- | --- | --- |
| weeks<br>23-25 | ‘pm’ | 0.49 | ambiguous<br>divergence |
| weeks<br>26-31 | ‘pm’ | 0.66 | moderate<br>similarity |
| weeks<br>32-34 | ‘pm’ | 0.27 | moderate<br>divergence |

**Table 3:**
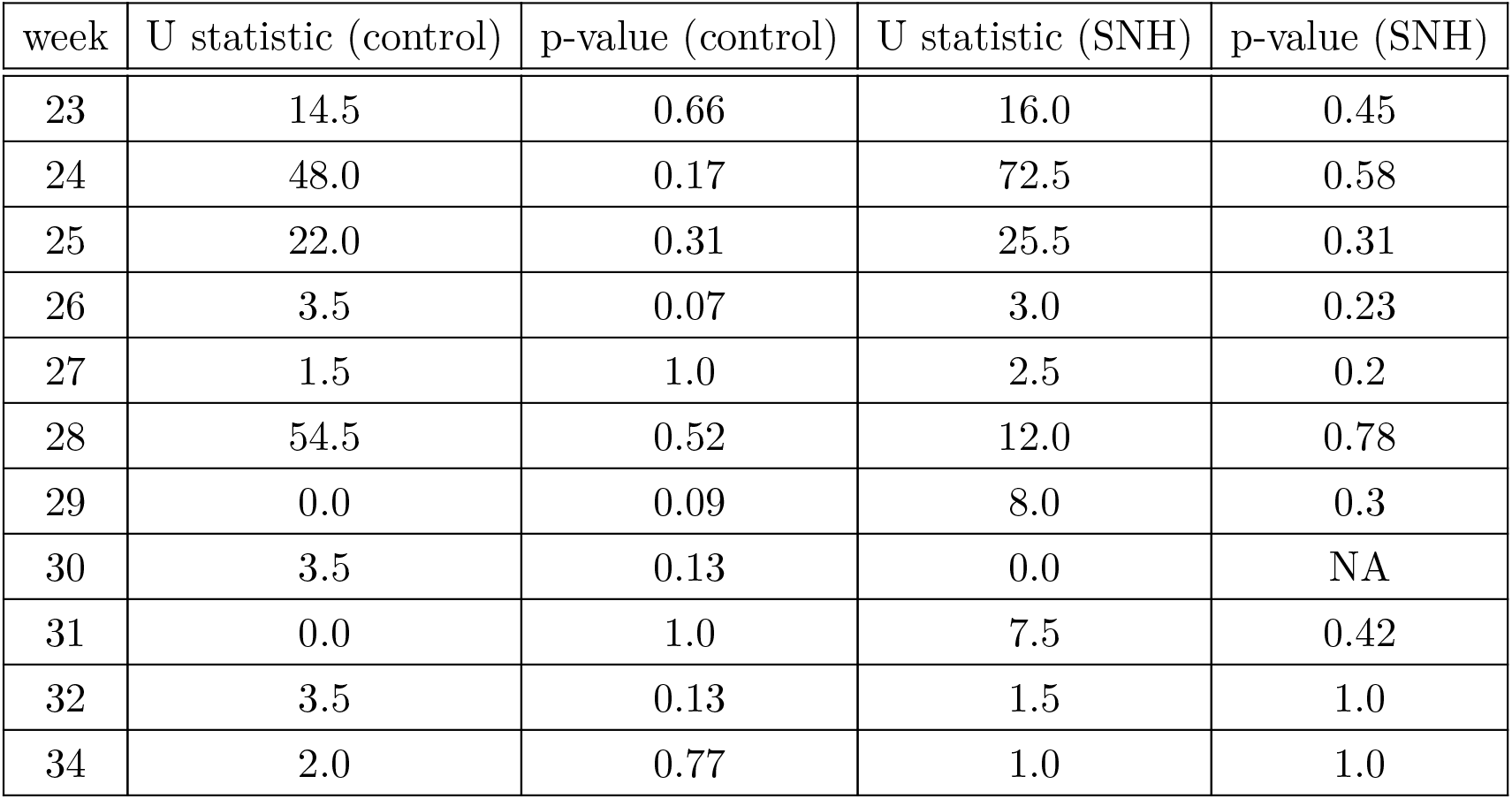
Intra-transect count similarity by week trap position 1-3 counts compared to trap position 8-10 counts daytime=‘pm’ Mann-Whitney U Test (Wilcoxon Rank Sum Test)

### 4.2 SNH vs. control transect comparison

#### 4.2.1 Inter-transect similarity by week

Neither morning nor afternoon data suggested a difference in weekly Thomisidae population between transects. The Mann-Whitney U test indicates that the count data does not contain enough evidence to conclude that the data comes from two different distributions. (minimum p-value = 0.16)

Bayesian estimation of the posterior distributions checked this result, the credible interval overlap coefficient used as as a similarity indicator.

The ‘overlap coefficient’ (Weitzman 1970; Pianka 1973; Hurlbert 1978) is designed to assess the similarity of two distributions, two posteriors in this case. A coefficient value of ‘0’ means ‘no similarity’, a value of ‘1’ would mean ‘exact similarity’.

bayestestR::overlap (Makowski, Ben-Shachar, and Lüdecke 2019) computes an overlap coefficient of the SNH and control posteriors as 0.49 in the first 3 weeks of the study, and 0.66 in the second weekly cluster. I assess 0.49 as ‘ambiguous similarity’ meaning that the two transects seem to share equivalent proportions of equivalent and different observed trap counts. A coefficient of 0.66 suggests to me that there is ‘moderate similarity’ of the counts in the second weekly cluster.

In this study, the credible interval (Zhao et al. 2023) refers to the uncertainty regarding trap counts (a parameter) while a confidence interval would refer to the null hypothesis that the observed distributions are equivalent. The shared span of the transect credible intervals in the second weekly cluster reinforces the assessment of at least ‘moderate similarity’ of the distributions.

Re-sampling methods assessed the overlap of the posterior distribution credible intervals as being generally ambiguous (Table 5).

**Table 4:** Thomisidae count data SNH vs control by week for daytime = ‘am’ and ‘pm’ Mann-Whitney U Test (Wilcoxon Rank Sum Test)

| week | U statistic (am) | p-value (am) | U statistic (pm) | p-value (pm) |
| --- | --- | --- | --- | --- |
| 23 | 60.5 | 0.42 | 60.5 | 0.25 |
| 24 | 36.5 | 0.32 | 36.5 | 0.40 |
| 25 | 35.5 | 0.27 | 35.5 | 0.30 |
| 26 | 36.5 | 0.29 | 36.5 | 0.41 |
| 27 | 42.5 | 0.54 | 42.5 | 0.47 |
| 28 | 35.5 | 0.39 | 38.5 | 0.79 |
| 29 | 64.5 | 0.26 | 64.5 | 0.22 |
| 30 | 55.0 | 0.71 | 55.0 | 0.18 |
| 31 | 67.5 | 0.16 | 67.5 | 0.43 |
| 32 | 60.0 | 0.37 | 60.0 | 0.47 |
| 34 | 55.0 | 0.37 | 55.0 | 0.30 |

**Table 5:** Similaarity assessment of SNH and control transect posterior count distributions.

| temporal cluster | sampling time | Overlap Coefficient | similarity assessment |
| --- | --- | --- | --- |
| weeks 23-25 | ‘am’ | 0.46 | ‘ambiguous divergence’ |
| weeks 26-30 | ‘am’ | 0.42 | ‘ambiguous divergence’ |
| weeks 31-34 | ‘am’ | 0.28 | divergence |
| weeks 23-25 | ‘pm’ | 0.49 | ‘ambiguous divergence’ |
| weeks 26-30 | ‘pm’ | 0.65 | ‘moderate similarity’ |
| weeks 31-34 | ‘pm’ | 0.27 | ‘moderate divergence’ |

### 4.3 The dataset profile

#### 4.3.1 Samples were zero-inflated

#### 4.3.2 Irregular spatial population distributions observed

### 4.4 Assessing the sampling design

Each transect sample comes from a triplet of vineyard rows. A theoretically uniform Thomisidae dispersion pattern and theoretically uniform trapping efficiency would make these triplets **ipso facto** identical.

I used the Wilcoxcon Signed Rank Test (‘SRT’) (Laureysens et al. 2004; Knight, Mahony, and Green 2014) to compare pairs of row count distributions. The 3 rows of an individual same-transect, same-time sample results in a similarity assessment between row 1 and row 2, row 1 and row 3, and row 2 and row 3. A three day sampling week for that transect/time context then results in 9 SRT comparisons (with associated p-values represented by the single boxplot).

Non-zero counts became sparse at the end of the sampling period making SRT comparison impossible in some cases.

The p-value boxplots indicate that there is not enough evidence to conclude that the sample count medians for the vineyard same-transect, same-time row pairs are different. The result gives me some confidence that the sampling design allowed me minimize bias related to the geometry of the transect in that traps were sufficiently spaced laterally to justify a claim of independence while minimizing the effect of local ecological (SNH canopy or vineyard) variation.

### 4.5 Row triplet pattern similarity

The surprising variation of Thomisidae counts obtained from triplets of same transect/day/ time sampled rows (Figure 7) was not anticipated by the vineyard research that I had reviewed. I had imagined my sampling design would expose a population transition gradient that might be explained primarily by the influence of SNH. I now wonder if effects based on Thomisidae ecology and/or the effect of blue vane traps may need to be evaluated.

#### Presence-absence data

Earlier vineyard studies mention Sørensen as a means of evaluating pairwise similarity, but, applied to the data of this study, the index does not seem informative and offers suspicious results both for inter-transect and intra-transect comparison.

There are at least four Sørensen Index variations: Sørensen Classic for presence/absence data, Sørensen for Binary Data (True Positives, False Positives/Negatives), Sørensen Quantitative (Czekanowski Index), and Sørensen Continuous. Generally, they are all approaching basic set theory comparison, doubling the intersection and dividing by the sum of set sizes. The variations adapt to data types (binary, continuous, abundance) and study designs (pairwise vs. multi-site). For example, the Czekanowski index generalizes Sørensen to abundance data, while the continuous version handles probabilistic outputs.

Set theory similarity measures in this context seem forced for convenience as transformation of counts into into a binary representation reduces the information contained in the sample and may obscure information about natural processes.

In this study, I had hoped to identify some suggestion of a change in population gradient. This objective implied that Sørensen Classic would be less useful. However, I noticed that the

Sørensen - Czekanowski Index applied to assess the similarity of same-transect observations seemed to lack a degree of sensitivity as there was little variation in the computed index. This transect data is not necessarily over-dispersed, but it is zero-inflated (Lee 2025), Figure 10.

**Figure 9.**
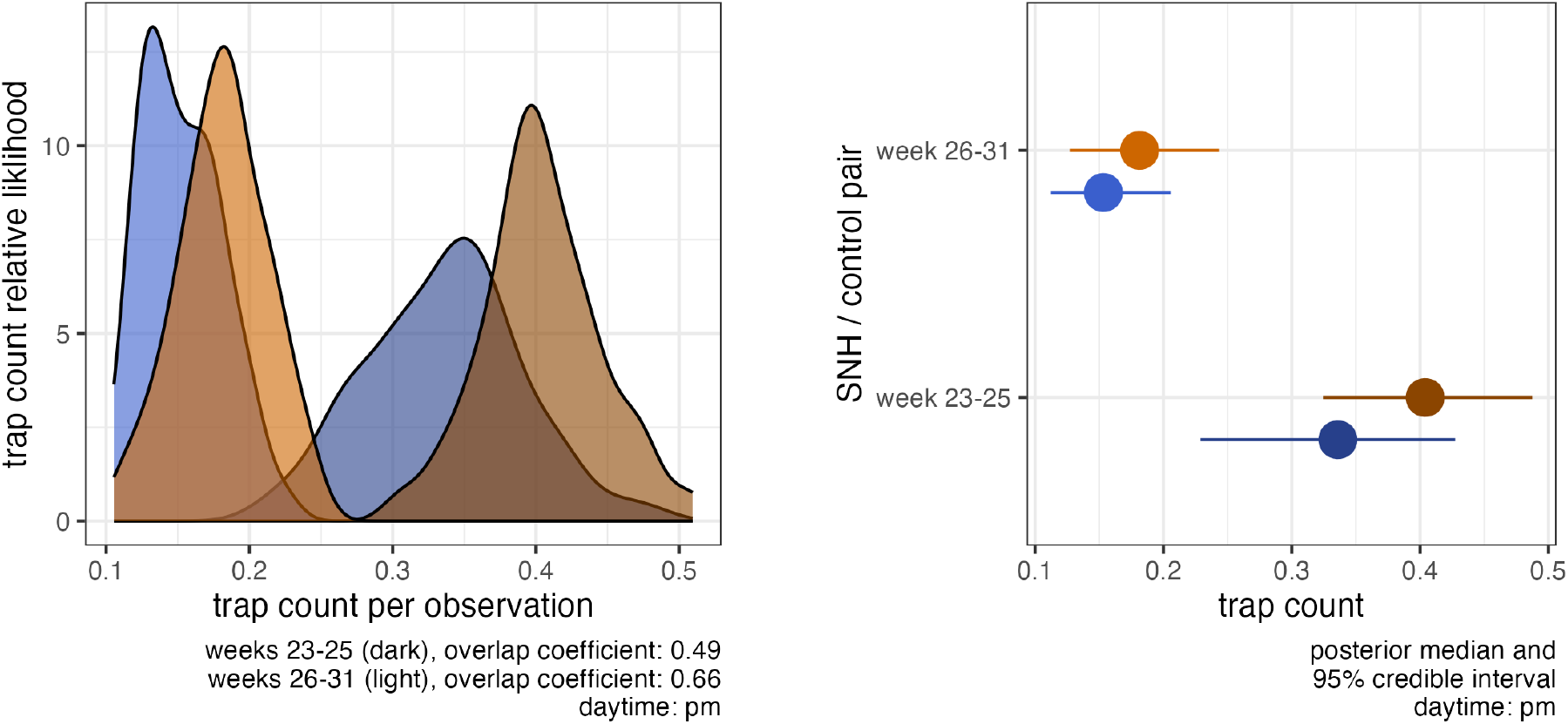
Simulated posterior distributions of Thomisidae counts weeks 23-25 and weeks 26-31.

**Figure 10.**
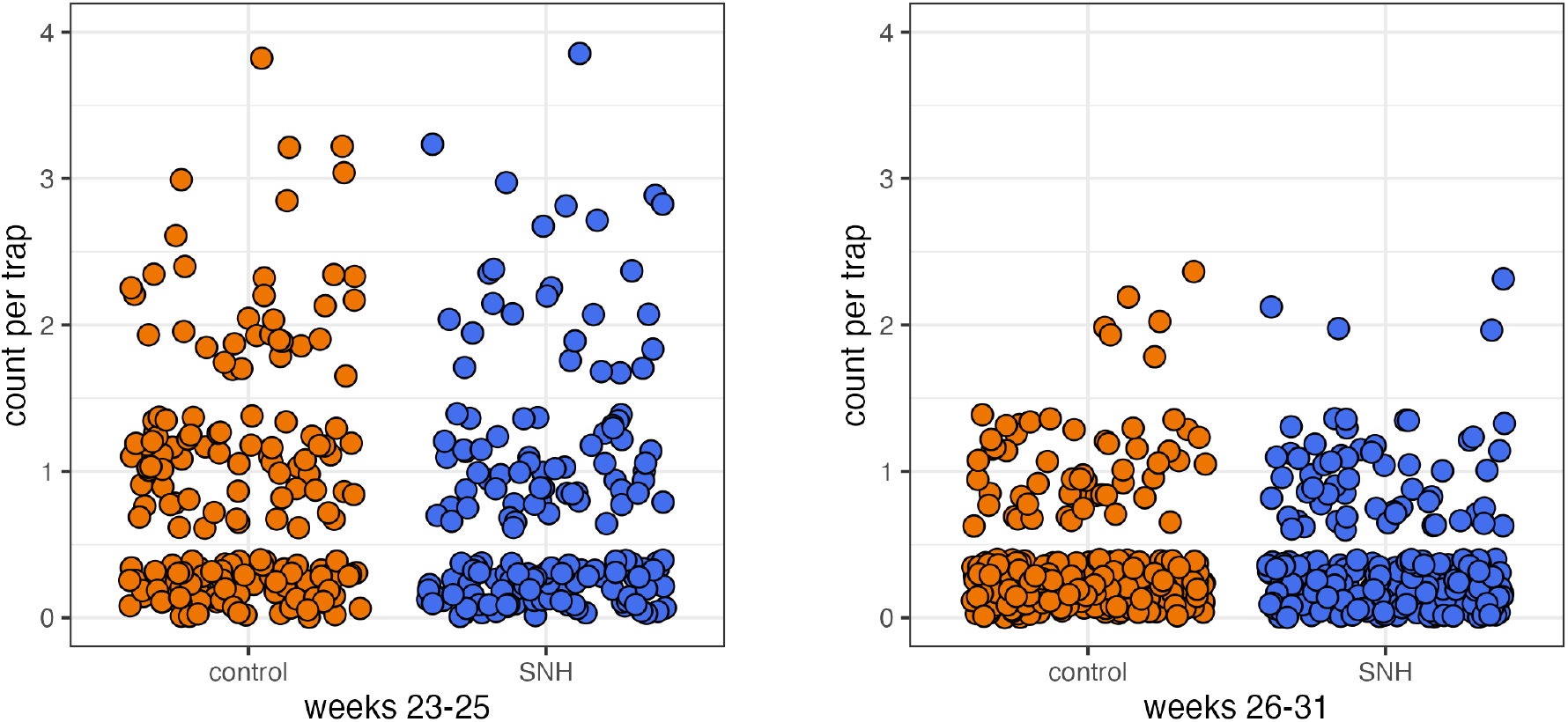
Thomisidae observations in two seasonal periods.

**Figure 11.**
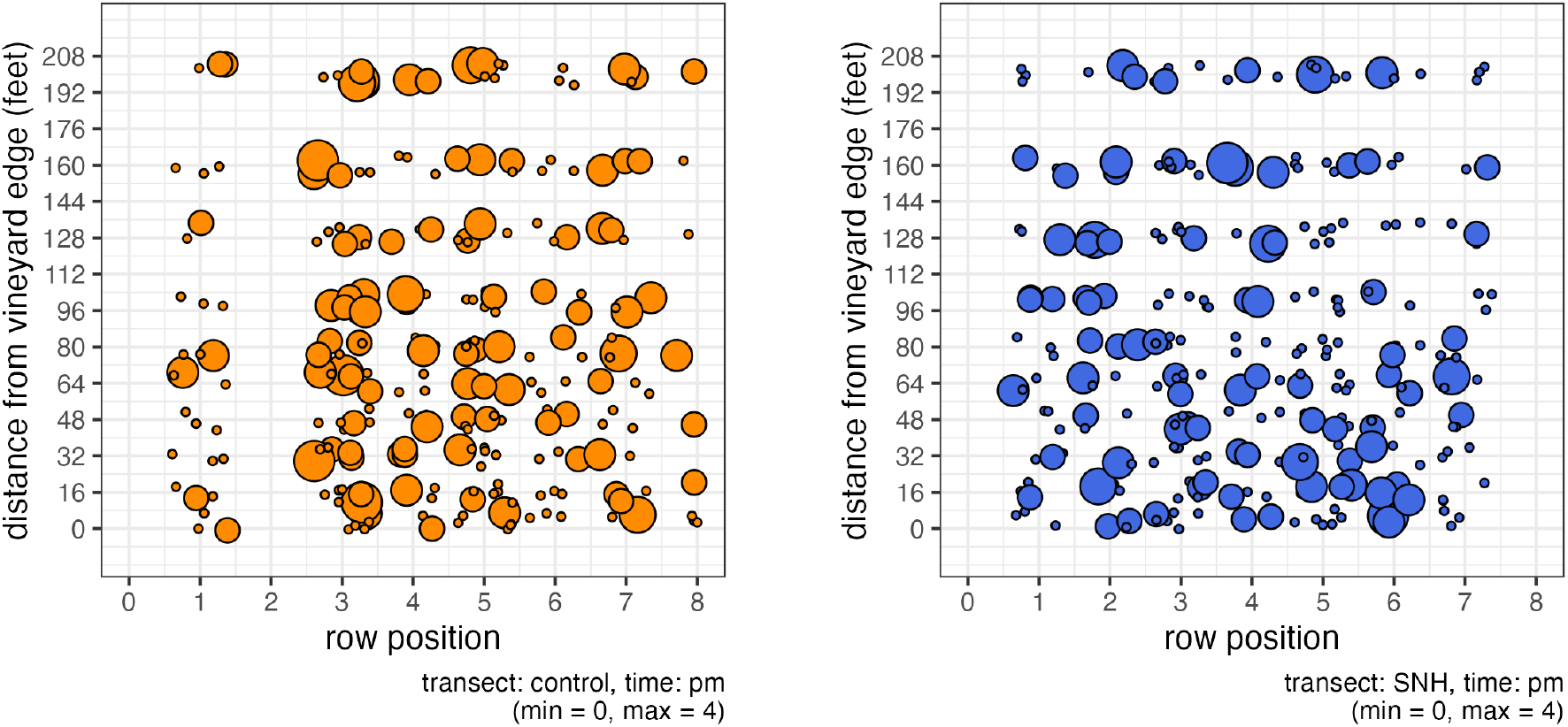
Thomisidae presence: distance by vineyard row position.

Experimenting with text that mimics presence-absence patterns of 3 ten trap vineyard rows, I adapted large language model (LLM) text comparison tools to evaluate their ability to assess the differences in presence-absence patterns.

LLMs use ‘N-Grams’ to capture word sequences by using a fixed character width, the vector representation of which then includes information about adjacent words, thus incorporating word order (Kondrak 2005).

The python library ngram includes an NGram.compare function (Lau 2013) that compares two strings (or sentences) based on a string (or ‘character’) length parameter and returns a score. For example,

~~~
#code(lang: “python”)
import ngram
similarity_score = ngram.NGram.compare(‘Ham’, ‘Spam’, N=1)
~~~

This text comparison results in a similarity score of 0.4

One option for converting presence-absence data for a 10 trap vineyard row into a string would be to use the character ‘T’ to indicate presence and the character ‘f’ to indicate absence:

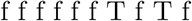

(I’m using typographic case and the insertion of spaces to make this pattern readable.) ngram.NGram.compare() can then compare this string to another string given a character width (the N-Gram) to drive the computation.

I can now take Thomisidae count data for each row sampled, convert to this presence-absence representation, and generate a similarity ‘score’ or ‘index’ between row pairs. Again, for a 3 row sampling sequence, the pairs would be row1/row2, row1/row3, and row2/row3, resulting in 3 N-Gram scores for that sampling event. Phase 1 of this data structure is represented by Table 6

**Table 6:**
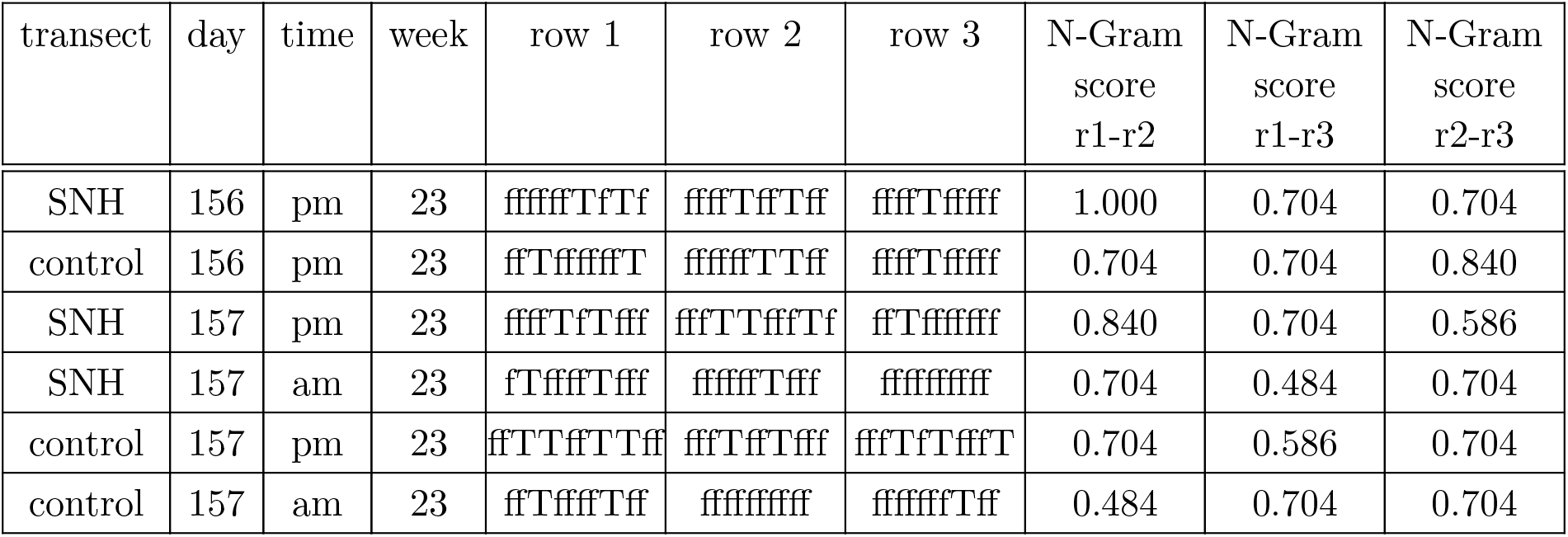
Example presence-absence data structure: phase 1.

Similarly, I can use the same 3 rows of presence-absence data to generate a Sørensen Index triplet. Phase 2 results when I add those to the data structure.

Now, I can evaluate the sensitivity of these two measures by generating 10,000 random presence-absence triplets and the related N-Gram and Sørensen indexes as shown in Figure 13. actually, this is not correct: the data is binary Czek expects counts.

**Figure 12.**
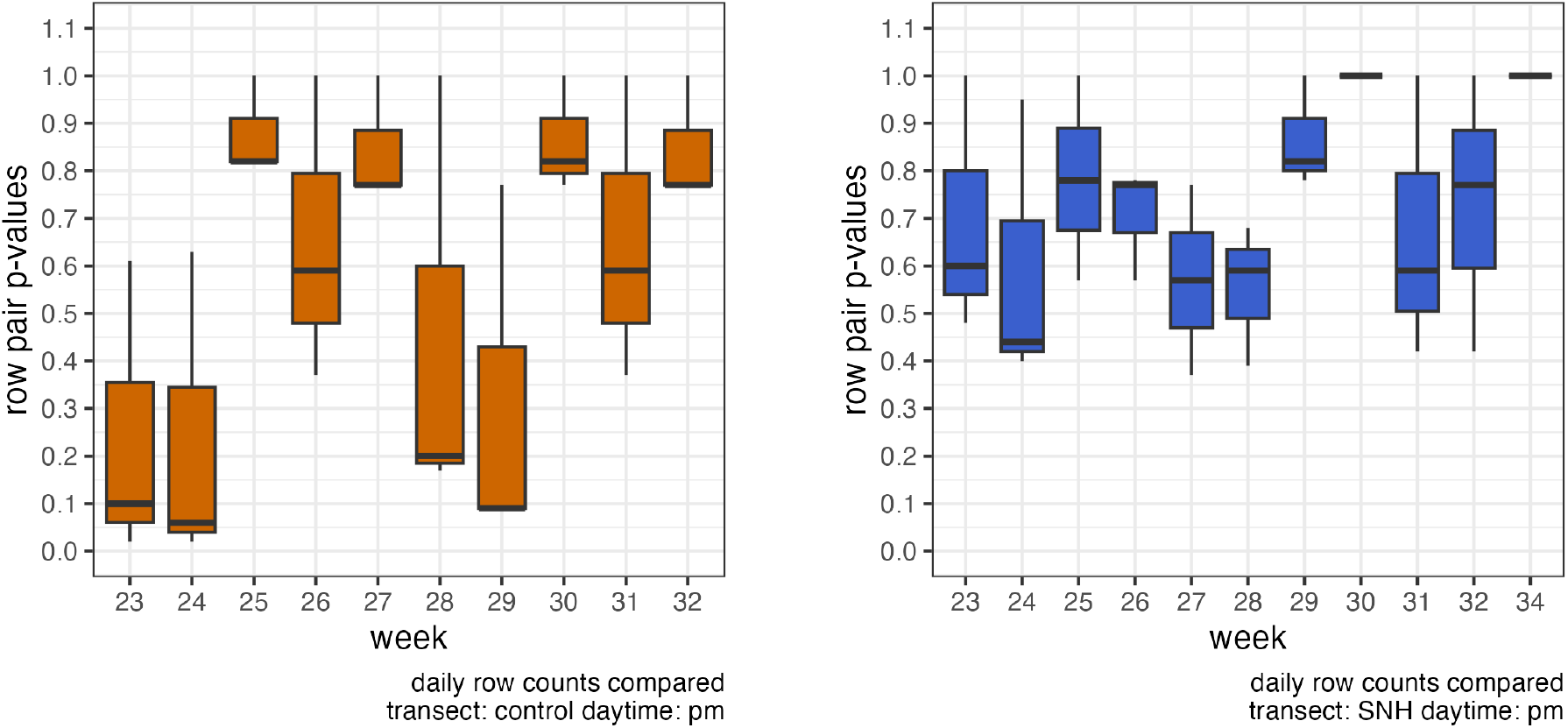
Thomisidae counts: same-transect, same-day comparison with Wilcoxcon SRT (daytime=‘pm’)

**Figure 13.**
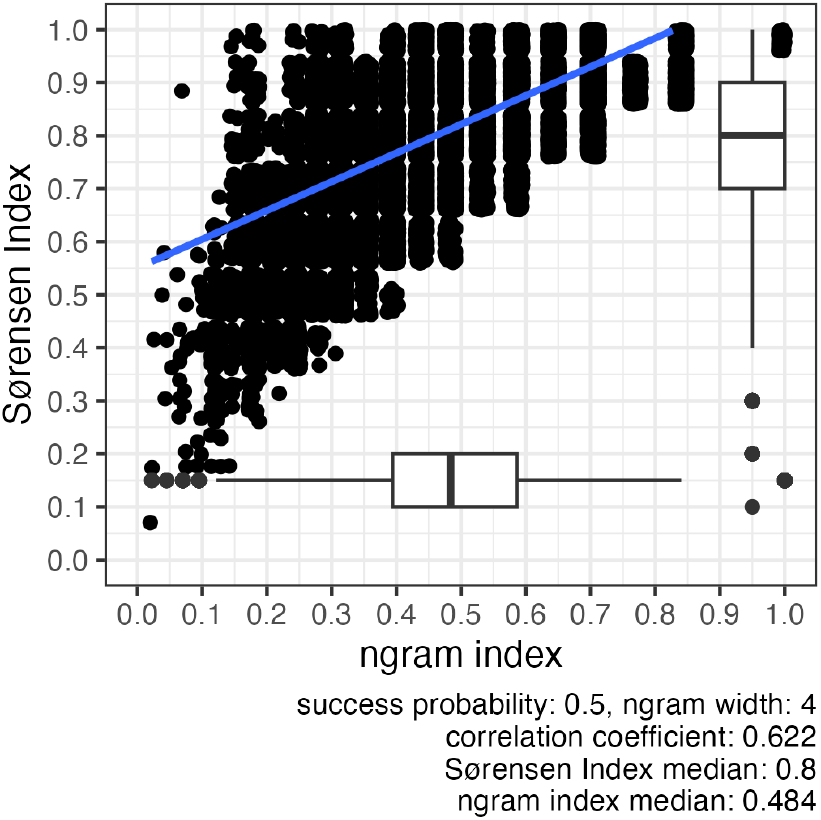
N-Gram vs. Sørensen-Czekanowski Index trap presence probability = 0.5.

I notice that with an equal probability (50%) of any of the 10 character positions being set to ‘T’, and the sliding N-Gram window being set to 4 characters, that the N-Gram index does, in fact, almost center on 0.5, a similarity score that an LLM model would possibly characterize as suggesting weak or perhaps moderate similarity. However, the Sørensen Index, processing the same data without any ability to recognize patterns, reports that these random counts are very similar.

Setting the Thomisidae trap success probability to 30% (per Figure 9) the NGRAM Index median increases slightly and the Sørensen Index median jumps to 0.9 (Figure 14).

**Figure 14.**
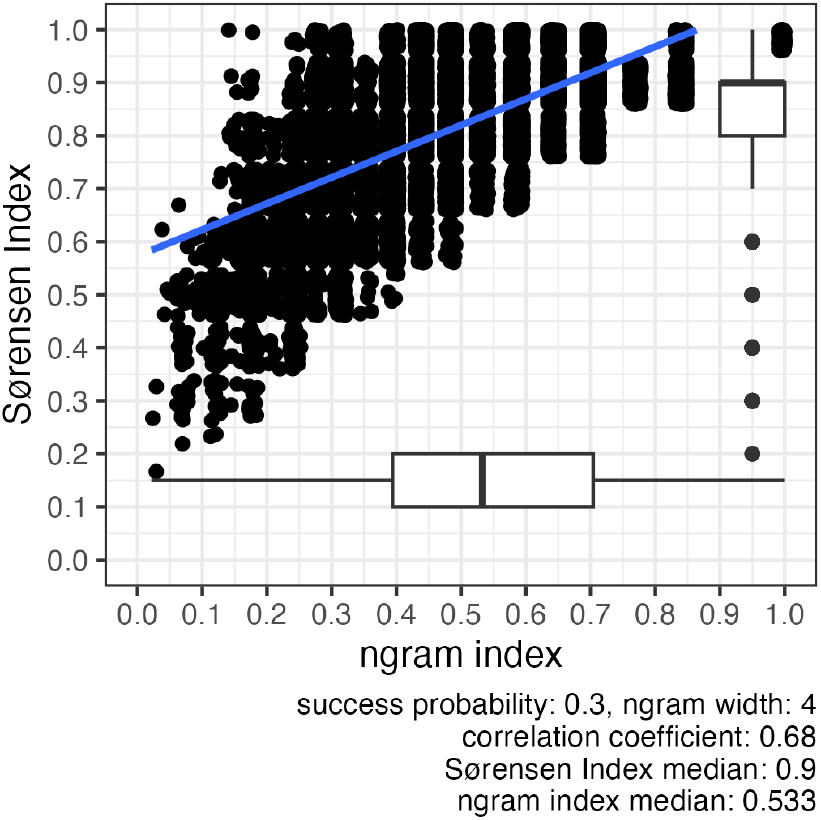
N-Gram vs. Sørensen-Czekanowski Index trap presence probability = 0.3.

My instinct is that the vine row association patterns of Figure 11 means that the assumption of single vine row trap count independence is questionable and that a presence-absence application is not a good fit for Sørensen. Perhaps this confounding effect of count patterns or bubbles made it impossible for earlier researchers to clearly identify SNH distance effects in their data.

## 4.6 Limitations

I do not have a sufficient understanding of the crab spider lifecycle in the oak forest and in the vineyard. Ideally, a subsequent study would include an assessment of the population in the oak habitat and the survey would start at least six weeks earlier.

Effects of vineyard operations were not assessed in this study, yet they may have influenced arthropod populations. Shoot thinning and leaf pulling intentionally change the canopy and the effect of the application of sulphur and horticultural oils were not evaluated.

### 4.7 Implications and Recommendations

Habitat profiles are an important characteristic when considering vineyard ecology. In the Central Coast, limited irrigation water availability makes cover crops difficult to maintain, ultimately influencing populations of arthropod predators and prey. High level SNH effects described by other studies may be unique to a specific ecology.

Central Coast SNH assessment projects could attempt to duplicate the collection strategies used in earlier studies to identify a broader range of vineyard insects and perhaps be more precise about SNH effects.

The blue vane trap layout and sampling strategy did not identify the hypothesized gradual change in Thomisidae density as the population dispersed relative to the SNH edge. It was effective, however, in collecting Diptera (Figure 3). Rather that focusing on beneficials, the blue vane traps could be used by grape growers to raise their awareness of pest density.

## 5 Conclusions

This study was unable to determine if the classic, semi-natural habitat associated with Central Coast vineyards actually promotes populations of the beneficial arthropods. Additionally, analysis of Thomisidae count data could not identify a population gradient extending from the SNH vineyard edge.

Despite the seemingly negative result, I am reminded that cover crop contributions are emphasized by prior SNH research. Central Coast vineyardists actively suppress native interrow cover, yet these plants may be the determinant of SNH effects.

This study did demonstrate that blue vane traps have some utility in monitoring arthropods in a vineyard canopy. Grape growers with moderate entomology aptitude can identify Thomisidae, an antagonist of common vineyard pests, using simple, non-destructive tools.

Also, the novel application of N-Gram pattern analysis in an ecology context may suggest that statistical research into N-Gram based analysis could reinforce traditional modelling.

My thought is the contribution of vineyard SNH might be more effectively expressed via positive economic effects that result from improved farm worker diligence at the operational level. I would encourage a study of the benefit of raising farm workers awareness by requiring them to monitor vane traps and to log observations while concurrently assessing the quality of canopy management and material application.

## Acknowledgments

I thank Peter Work, Ampelos Cellars, for hosting this study in his vineyard.

I thank Dr. Lindsey Norgrove, Bern University of Applied Sciences, for her help in focusing the research question and formalizing a literature review..

I’m grateful to Ric Fuller, Viticulture Instructor, Allan Hancock College for his encouragement and numerous insights regarding beneficials and sampling.

## Appendix A

### Arthropods sampled using blue vane traps and their ranked abundance

**Table 7:**
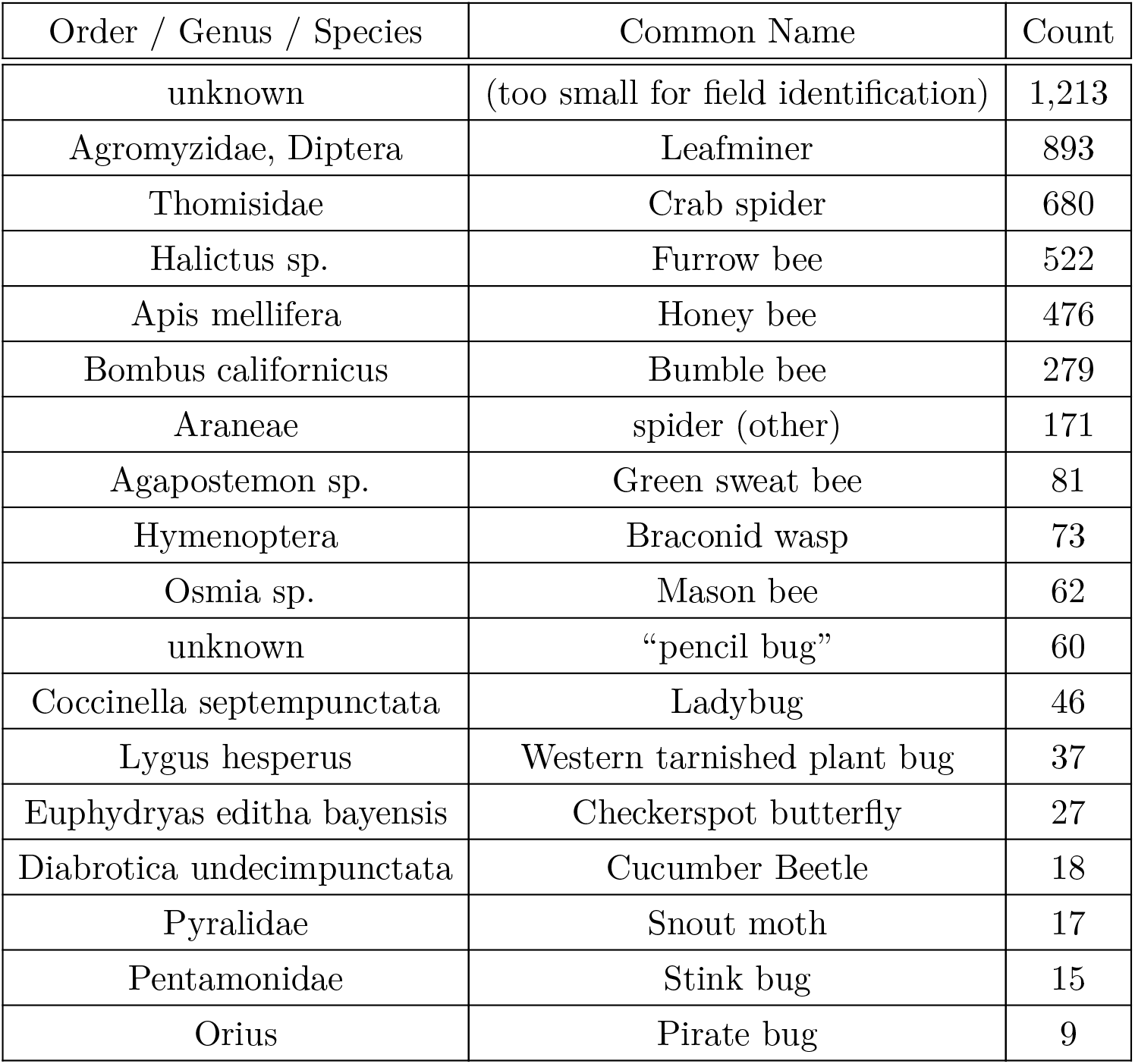
arthropods sampled.

## Appendix B

### Sta. Rita Hills AVA 2018 weather conditions

The California Department of Water Resources maintains the California Irrigation Management Information System (CIMIS) network of weather stations that serve weather data to the public. (Water Resources 2018)

The CIMIS 231 weather station (latitude 34.672222, longitude −120.51306) is approximately 7.71 km (4.79 miles) from the Ampelos vineyard.

**Figure 15.**
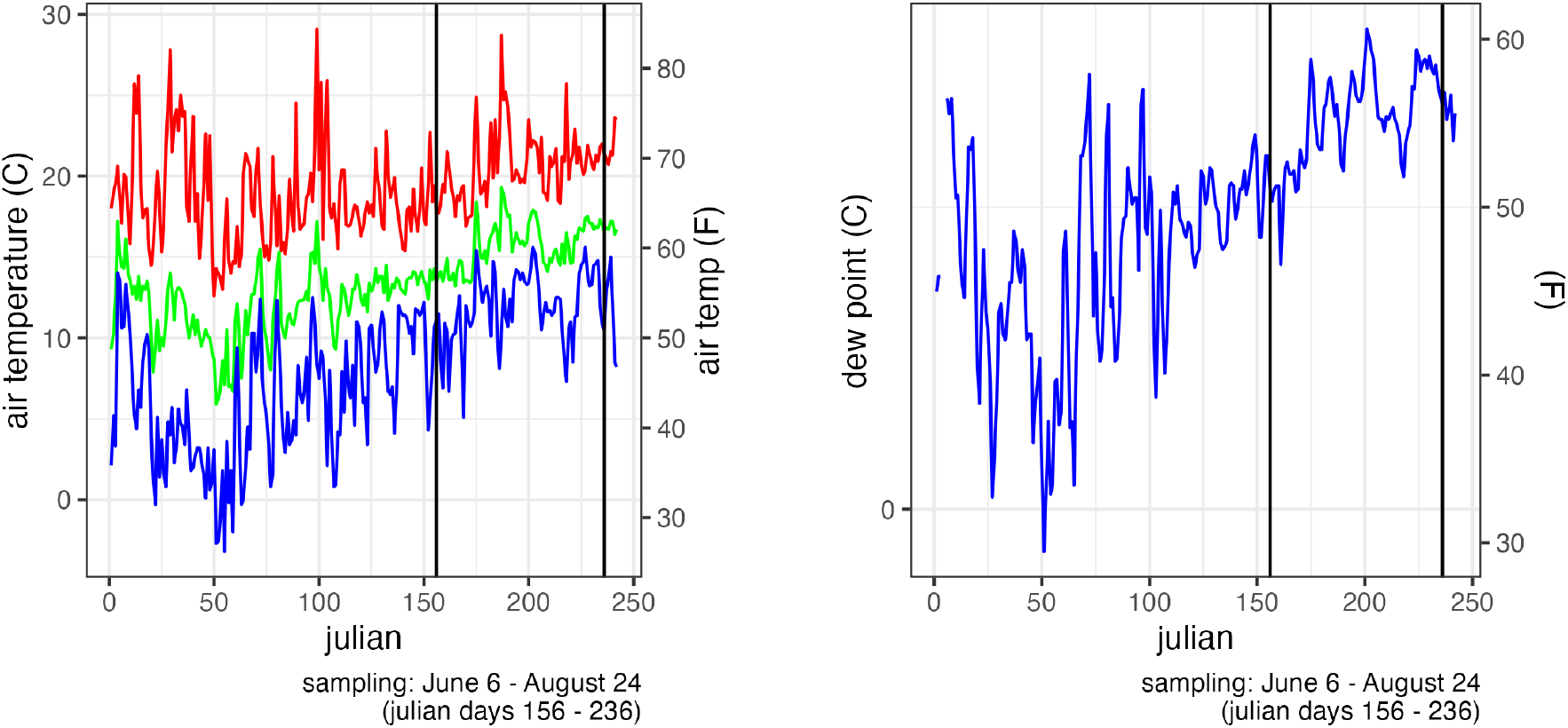
2018 CIMIS 231 air temperature and dew point data.

**Figure 16.**
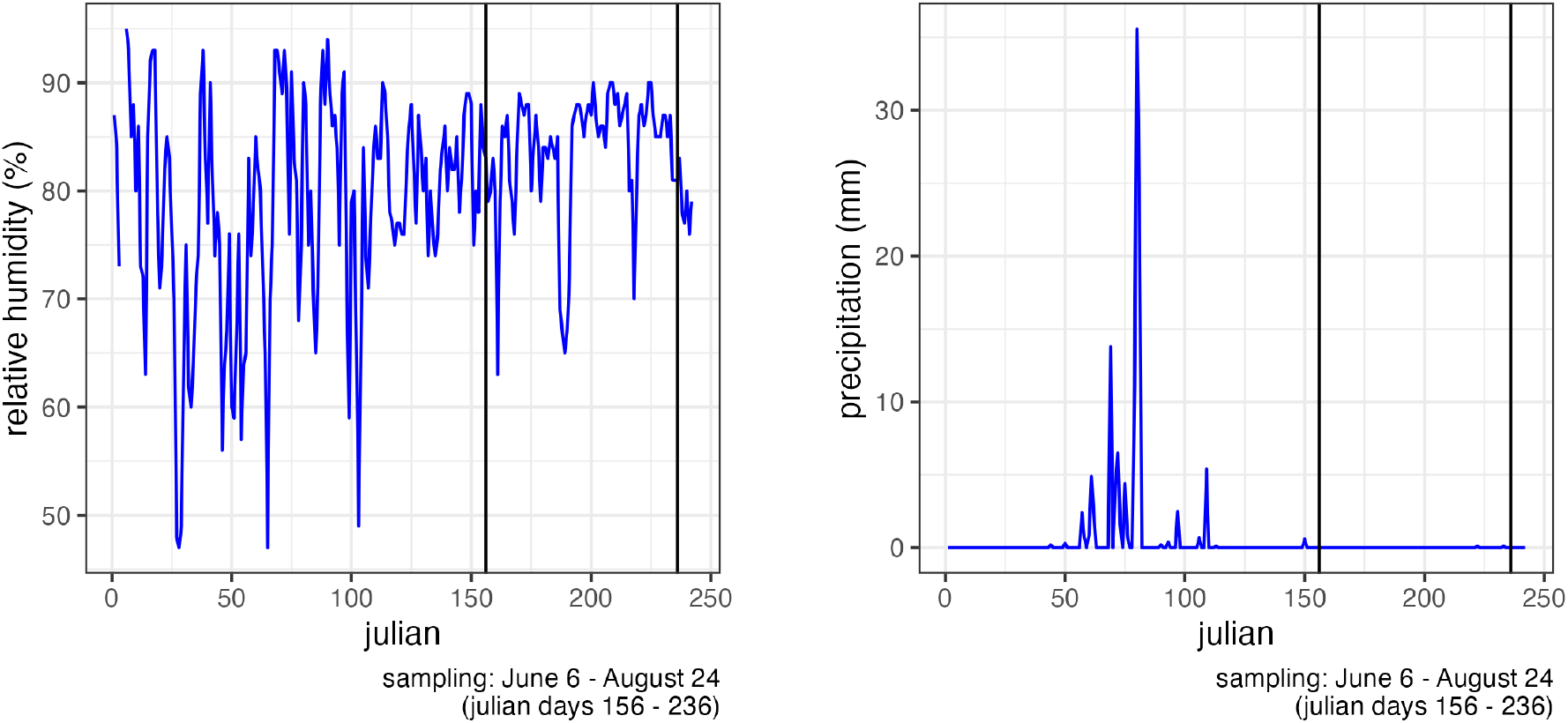
2018 CIMIS 231 humidity and precipitation data.

**Figure 17.**
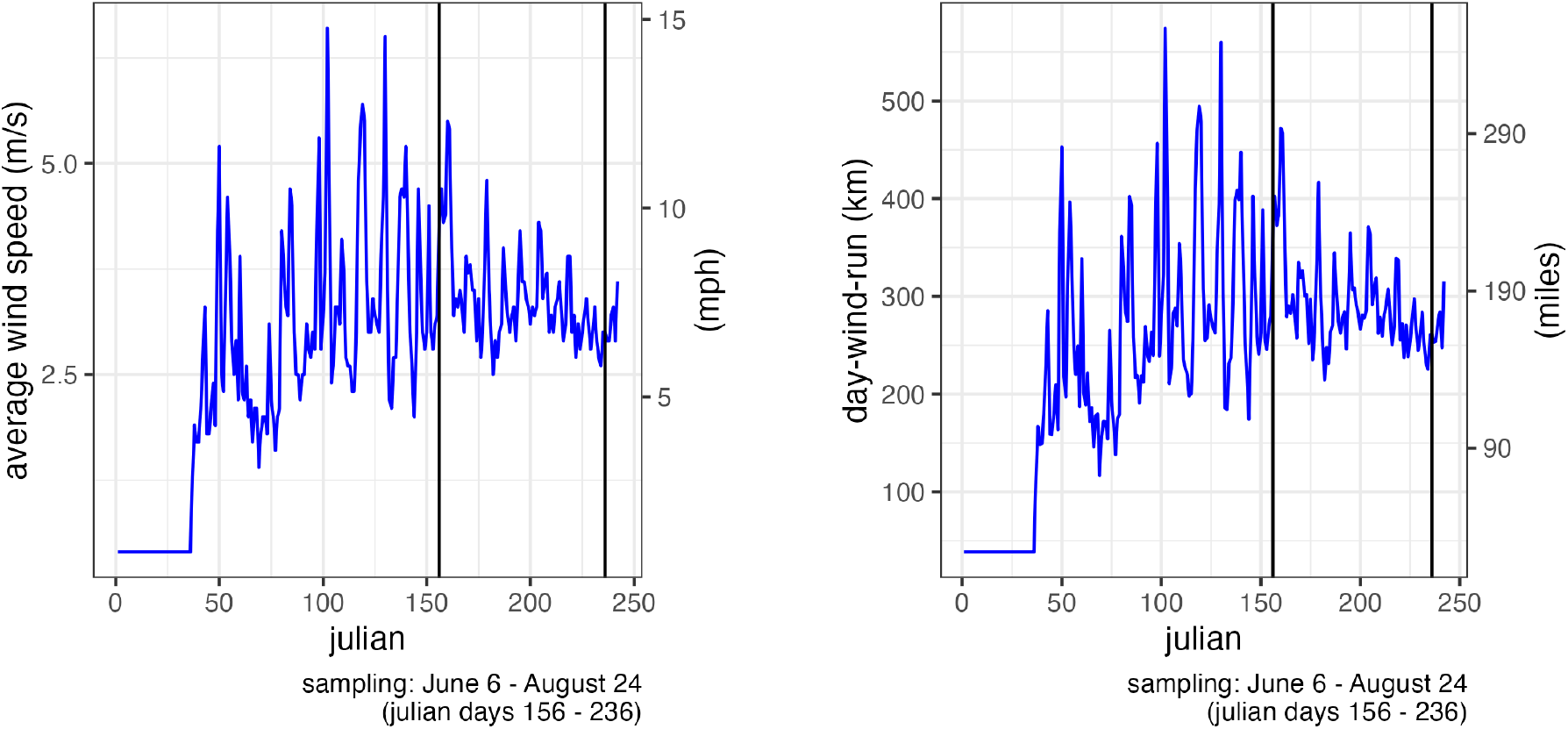
2018 CIMIS 231 wind speed and wind run data.

## Appendix C

### access to the study data set

~~~
#code(lang: “R”)
source.url = c(“https://raw.githubusercontent.com/cordphelps/ampelos/master/data/bugs.csv“)
bugs.tibl = read.csv(source.url, header=TRUE, row.names=NULL)
or
#code(lang: “python”)
import pandas as pd
import requests
import csv
import io
result_list = []
url = ‘https://raw.githubusercontent.com/cordphelps/ampelos/refs/heads/master/data/bugs.csv’response = requests.get(url)
# Check if the request was successful
if response.status_code == 200:
  csv_data = response.text
  # Use csv.reader to read the CSV data
  # Use io.StringIO to convert the bytes to a string records = []
  reader = csv.reader(io.StringIO(csv_data), delimiter=‘,’) for row in reader: if row: # Ensure the row is not empty records.append(row) result_list = records
# pandas: list to df
bugs_df = pd.DataFrame(result_list)
~~~

## Appendix D

### software used to produce the data analysis

**Table 8:**
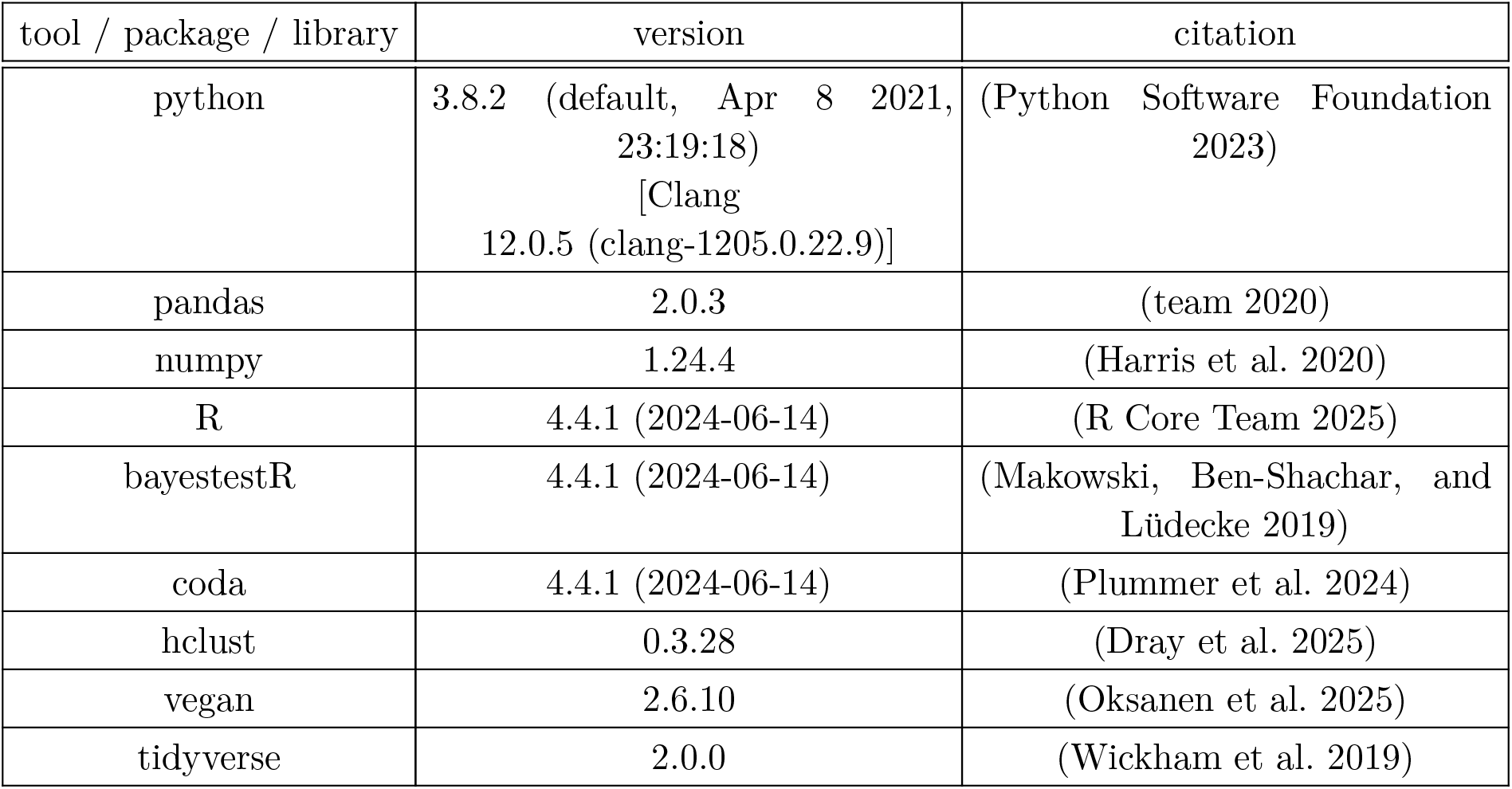
3rd party software versions.

## Notes

https://github.com/cordphelps/ampelos

https://github.com/cordphelps/ampelos-similarity

### Competing Interest Statement

The authors have declared no competing interest.

### Summary of Updates

grammatical corrections in the heading of Appendix C

## References

Supporting project material may be found on https://github.com/cordphelps/ampelos

Custom software supporting the project analysis may be found on https://github.com/cordphelps/ampelos-similarity

## Bibliography

[1] M. A. Altieri, C. I. Nicholls, H. Wilson, and A. Miles, Habitat Management in Vineyards: A Growers Manual for Enhancing Natural Enemies of Pests. Berkeley, CA, USA: Laboratory of Agroecology, College of Natural Resources, University of California, 2010. [Online]. Available: https://agroecology.berkeley.edu/resources/Altieri_2010_habitat_management_in_vineyards.pdf

[2] BanfieldBio, “Blue Vane Traps.” 2025.

[3] B. Valentini, F. Barbero, L. P. Casacci, A. Luganini, and I. Stefanini, “Forests Influence Yeast Populations Vectored by Insects into Vineyards,” Frontiers in Microbiology, vol. 12, p. 1039939, 2022, doi: 10.3389/fmicb.2022.1039939.

[4] D. G. James, S. Link, and T. L. Pyle, “Manipulating Vineyard Biodiversity for Improved Insect Pest Management: Final Report for SARE Project SW10-052,” 2013. [Online]. Available: https://projects.sare.org/project-reports/sw10-052/

[5] C. I. Nicholls, M. Parrella, and M. A. Altieri, “The effects of a vegetational corridor on the abundance and dispersal of insect biodiversity within a northern California organic vineyard,” Landscape Ecology, vol. 16, no. 2, pp. 133–146, 2001, doi: 10.1023/A:1011117107468.

[6] N. E. Seavy, S. Quader, J. D. Alexander, and C. J. Ralph, “Considerations for the Design and Analysis of Monitoring Studies,” in Bird Conservation Implementation and Integration in the Americas: Proceedings of the Third International Partners in Flight Conference, C. J. Ralph and T. D. Rich, Eds., in General Technical Report PSW-GTR-191. Albany, CA, 2005, pp. 744–753. [Online]. Available: https://www.fs.usda.gov/psw/publications/documents/psw_gtr191/Asilomar/pdfs/744-753.pdf

[7] D. Makowski, M. S. Ben-Shachar, and D. Lüdecke, “bayestestR: Describing Effects and their Uncertainty, Existence and Significance within the Bayesian Framework,” Journal of Open Source Software, vol. 4, no. 40, p. 1541, 2019, doi: 10.21105/joss.01541.

[8] A. D. S. Inácio and M. X. Rodríguez-Álvarez, “Bayesian nonparametric inference for the overlap coefficient,” Statistical Methods in Medical Research, vol. 32, no. 1, pp. 136–151, 2023, doi: 10.1177/09622802221125573.

[9] B. N. Hogg and K. M. Daane, “The role of dispersal from natural habitat in determining spider abundance and diversity in California vineyards,” Agriculture, Ecosystems & Environment, vol. 141, no. 3–4, pp. 304–313, 2011, doi: 10.1016/j.agee.2011.03.003.

[10] Oregon Wine Research Institute, “Viticulture & Enology Technical Newsletter, Fall 2018.” 2018.

[11] R Core Team, “R: A Language and Environment for Statistical Computing.” Vienna, Austria, 2025. [Online]. Available: https://www.r-project.org/

[12] Y. Zhao et al., “The clustering of spatially associated species unravels patterns in tropical tree species distributions,” Ecosphere, vol. 14, no. 6, p. e4589, 2023, doi: 10.1002/ecs2.4589.

[13] C. Ricotta and J. Podani, “On some properties of the Bray-Curtis dissimilarity and their ecological meaning,” Ecological Complexity, vol. 31, pp. 201–205, 2017, doi: 10.1016/j.ecocom.2017.07.003.

[14] I. Zarraonaindia et al., “Soil management impacts vineyard soil fungal communities and their potential to influence grapevine health and wine quality,” Scientific Reports, vol. 8, no. 1, p. 11021, 2018, doi: 10.1038/s41598-018-29346-1.

[15] L. Mestre et al., “Both woody and herbaceous semi-natural habitats are essential for spider overwintering in European farmland,” Agriculture, Ecosystems & Environment, vol. 267, pp. 141–146, 2018, doi: 10.1016/j.agee.2018.08.027.

[16] M. A. Altieri, C. I. Nicholls, L. Ponti, and A. York, “Designing biodiverse, pest-resilient vineyards through habitat management,” Practical Winery & Vineyard Magazine, pp. 1–6, May 2005, [Online]. Available: https://agroecology.berkeley.edu/resources/Altieri_2005_Designing_biodiverse_pestresilient_vineyards.pdf

[17] M. J. Costello and K. M. Daane, “Abundance of spiders and insect predators on grapes in central California,” Journal of Arachnology, vol. 27, pp. 531–538, 1999.

[18] L. J. Thomson and A. A. Hoffmann, “Vegetation increases the abundance of natural enemies in vineyards,” Biological Control, vol. 49, no. 3, pp. 259–269, 2009, doi: 10.1016/j.biocontrol.2009.06.003.

[19] C. Kimoto, S. J. DeBano, R. W. Thorp, S. Rao, and W. P. Stephen, “Investigating temporal patterns of a native bee community in a remnant North American bunchgrass prairie using blue vane traps,” Journal of Insect Science, vol. 12, no. 108, pp. 1–23, 2012, doi: 10.1673/031.012.10801.

[20] M. Hall, “Blue and yellow vane traps differ in their sampling effectiveness for wild bees in both open and wooded habitats,” Agricultural and Forest Entomology, vol. 20, no. 4, pp. 487–495, 2018.

[21] M. J. Costello and K. M. Daane, “Day vs. Night Sampling for Spiders in Grape Vineyards,” The Journal of Arachnology, vol. 33, no. 1, pp. 25–32, 2005, doi: 10.1636/h02-52.

[22] J. M. Holland, J. N. Perry, and L. Winder, “The within-field spatial and temporal distribution of arthropods in winter wheat,” Bulletin of Entomological Research, vol. 89, no. 6, pp. 499–513, 1999, doi: 10.1017/S0007485399000653.

[23] S. Pearce and M. P. Zalucki, “Do predators aggregate in response to pest density in agroecosystems? Assessing within-field spatial patterns,” Journal of Applied Ecology, vol. 43, no. 1, pp. 128–140, 2006, doi: 10.1111/j.1365-2664.2006.01116.x.

[24] S. Dray et al., “adespatial: Multivariate Multiscale Spatial Analysis.” 2025. [Online]. Available: https://cran.r-project.org/package=adespatial

[25] G. Guénard and P. Legendre, “Hierarchical Clustering with Contiguity Constraint in R,” Journal of Statistical Software, vol. 103, no. 7, pp. 1–26, 2022, doi: 10.18637/jss.v103.i07.

[26] M. S. Weitzman, “Measures of overlap of income distributions of white and Negro families in the United States,” Washington, D.C., 1970.

[27] E. R. Pianka, “The structure of lizard communities,” Annual Review of Ecology and Systematics, vol. 4, pp. 53–74, 1973, doi: 10.1146/annurev.es.04.110173.000413.

[28] S. H. Hurlbert, “The measurement of niche overlap and some relatives,” Ecology, vol. 59, no. 1, pp. 67–77, 1978, doi: 10.2307/1936632.

[29] I. Laureysens, R. Blust, L. De Temmerman, C. Lemmens, and R. Ceulemans, “Clonal variation in heavy metal accumulation and biomass production in a poplar coppice culture: Seasonal variation in leaf, wood and bark concentrations,” Environmental Pollution, vol. 131, no. 3, pp. 485–494, 2004, doi: 10.1016/j.envpol.2004.02.009.

[30] E. C. Knight, N. A. Mahony, and D. J. Green, “Crop type influences edge effects on the reproduction of songbirds in sagebrush habitat near agriculture,” Avian Conservation and Ecology, vol. 9, no. 1, p. 8, 2014, doi: 10.5751/ACE-00662-090108.

[31] S. Lee, “Zero-Inflated Models 101: 10 Essential Facts for Data Scientists.” [Online]. Available: https://www.numberanalytics.com/blog/zero-inflated-models-essential-facts-data-science

[32] G. Kondrak, “N-Gram Similarity and Distance,” in String Processing and Information Retrieval (SPIRE 2005), in Lecture Notes in Computer Science, vol. 3772. Berlin, Heidelberg: Springer, 2005, pp. 115–126.

[33] J. K. S. Lau, “ngram: N-Gram String Similarity Module for Python.” 2013.

[34] C. D. of Water Resources, “CIMIS weather data, Station 231 (Glenbrook).” 2018.

[35] Python Software Foundation, “Python Language Reference, version 3.x.” 2023. [Online].Available: https://www.python.org/

[36] T. pandas development team, “pandas-dev/pandas: Pandas.” [Online]. Available: 10.5281/zenodo.3509134

[37] C. R. Harris et al., “Array programming with NumPy,” Nature, vol. 585, no. 7825, pp. 357–362, Sep. 2020, doi: 10.1038/s41586-020-2649-2.

[38] M. Plummer, N. Best, K. Cowles, and K. Vines, “coda: Output Analysis and Diagnostics for MCMC.” 2024. [Online]. Available: https://cran.r-project.org/package=coda

[39] J. Oksanen et al., “vegan: Community Ecology Package.” 2025. [Online]. Available: https://cran.r-project.org/package=vegan

[40] H. Wickham et al., “Welcome to the tidyverse,” J. Open Source Softw., vol. 4, no. 43, p. 1686, Nov. 2019, doi: 10.21105/joss.01686.

